# Efficacy of chronic ultrasound neurostimulation on behaviors and distributed brain metabolism in depressive-like mice

**DOI:** 10.1101/813006

**Authors:** Marc Legrand, Laurent Galineau, Anthony Novell, Barbara Planchez, Bruno Brizard, Samuel Leman, Clovis Tauber, Jean-Michel Escoffre, Antoine Lefèvre, Philippe Gosset, Wissam El-Hage, Patrick Emond, Catherine Belzung, Ayache Bouakaz

## Abstract

Major depression is one of the main factors contributing to the Global Burden of Disease. Current treatment strategies (*e.g.,* antidepressants and neurostimulation techniques) of major depression show some limitations including inaccuracy and invasiveness. Ultrasound neurostimulation (USNS) has been recently introduced as a physical non-invasive method for brain tissue stimulation and has gained increasing interest. In this study, we sought to evaluate the efficacy of transcranial USNS in an unpredictable chronic mild stress (UCMS) mouse model. The results show that transcranial USNS of the infralimbic cortex reduced anxiety-related behaviors as well as some, but not all, depression-related parameters. [^18^F]-FDG microPET imaging and brain metabolomic analyses showed that USNS triggered the activation of targeted brain region in addition to brain areas at a distance from the targeted zone, alleviating anxiety and depression-related behaviors induced by the UCMS regimen. Transcranial ultrasound neurostimulation show therapeutic potential in some aspects of major depression.

## Introduction

According to the World Health Organization, major depression (MD) has already become the second most prevalent cause of illness-induced disability (1), which makes this disorder one of the main contributors to the Global Burden of Disease (2). It is generally treated with chronic antidepressants (ADs), which consist of drugs increasing monoaminergic neurotransmission such as selective serotonin reuptake inhibitors (SSRIs). However, nearly 65% of the patients do not respond to this first-line therapy, and it is established that 30-50% of patients are resistant to AD compounds (3, 4), which means they do not show remission after treatment with several ADs: this condition is conventionally referred to as treatment resistant depression (TRD). Furthermore, MD often presents itself with a high and disabling comorbidity to anxiety disorders, where SSRIs show a less reliable spectrum of therapeutic efficacy (5).

In the past decade, progress in the treatment of TRD has involved neurostimulation, which consists in activating/inhibiting the cerebral networks whose functioning is modified in MD. Regions of interest include the dorsolateral prefrontal cortex (dlPFC), the subgenual part of the anterior cingulate cortex (sgACC), the nucleus accumbens or the lateral habenula (6). While the dlPFC is a cortical area that can be targeted using neurostimulation methods such as repeated transcranial magnetic stimulation (rTMS) or direct current stimulation (DCS), the same does not apply to deeper brain regions that can be targeted only using invasive approaches such as deep brain stimulation (DBS). Therefore, there is an urgent need to develop novel and efficient neurostimulation techniques that can target deep brain regions in a focal and non-invasive manner.

Ultrasound (US) technology has been evaluated as a therapeutic tool in neurology/psychiatry either to induce non-invasive surgical ablation of a given brain region (7–9), to potentiate drug delivery (10–12), or more recently, to induce neuromodulation of a specific brain area (13–16). Ultrasound stimulation offers the advantages of being focused and able to activate deep regions of the brain, which is quite relevant to treat psychiatric disorders. However, US neurostimulation (USNS) has yet to be evaluated for this indication.

In this context, the objective of this study was to assess whether repeated transcranial USNS of the infralimbic cortex (IL), i.e. the sgACC counterpart in rodents (17), can counteract modifications induced by the unpredictable chronic mild stress (UCMS) procedure in mice. The UCMS model is often considered a naturalistic model of MD, in that it satisfies some criteria for face, predictive, and construct validity (18, 19). The sgACC was selected as a relevant target region because a) its activity patterns has been shown to be modified in MD patients (20, 21), b) repeated DBS of this region elicited therapeutic effects in TRD patients (22), c) using a mouse model, we have observed in a previous study that UCMS elicited changes in gene expression in this region that were partly reversed by chronic treatment with fluoxetine, an SSRI (23), d) DBS of this region in mice induced therapeutic-like effects using the UCMS model (24).

Optimal US parameters were first assessed by evaluating the muscular responses to single-pulse USNS of the contralateral primary motor cortex. These parameters were then used to investigate the effects of acute repeated USNS targeted to the IL. Chronic repeated USNS of the IL was then evaluated as a treatment against classic fluoxetine using a 5-week UCMS model combined with behavioral analyses of anhedonia and anxiety, short-term [^18^F]-FDG microPET imaging to assess the cerebral networks engaged by the repeated stimulations and a metabolomic analysis of several regions of the corticolimbic network to explore longer-term related mechanisms.

## Results

### Determination of optimal US parameters through single-pulse USNS of the primary motor cortex

US waves were generated using a single-element transducer with a central frequency of 500 kHz, coupled to a water-filled collimated column (d=10 mm) and operated on a stereotaxic frame. The transducer had a diameter of 38 mm and was focused at 65 mm, driven with an electrical signal generated by an arbitrary waveform generator set for 160-millisecond pulses and amplified with a power amplifier. To determine optimal US parameters as a basis for repeated USNS treatments, several criteria (success of stimulation, strength of stimulation and reliability across subjects) were assessed with single-pulse USNS delivered over the left primary motor cortex M1 (bregma −0.5 mm). In a cohort of thirty mice positioned in the stereotaxic frame under gaseous anesthesia (0.1-1% isoflurane), peak negative pressures ranging from 50 to 500 kPa by steps of 50 kPa were delivered from the transducer (25) (Figure 1A).

**Figure 1:**
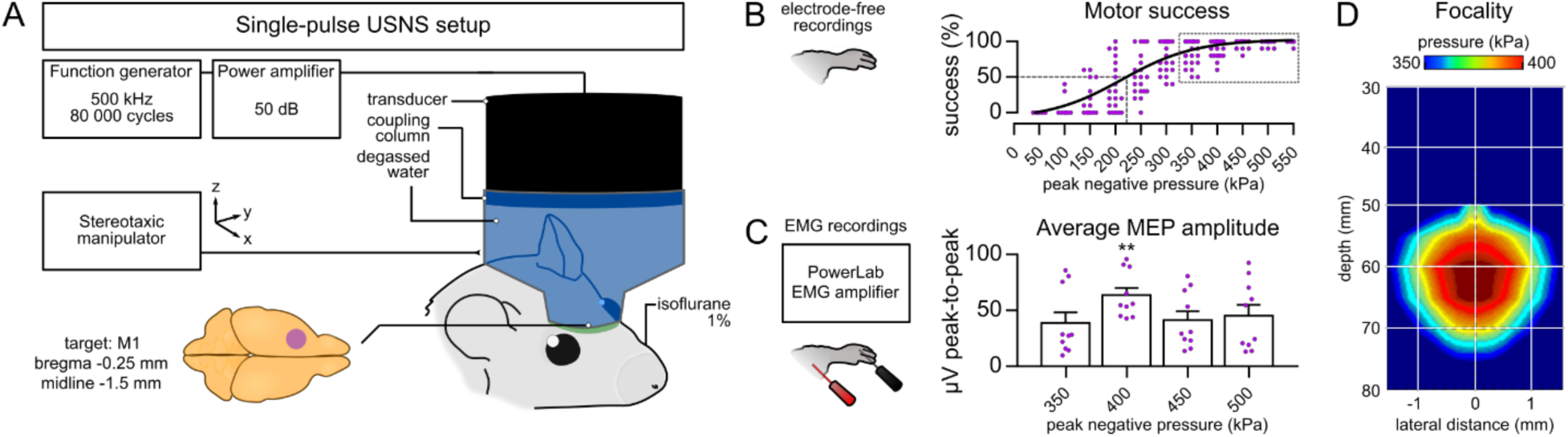
Determination of optimal US parameters. (A) Schematic view of the system for single-pulse USNS and brain targeting of primary motor cortex M1. (B) Electrode-free recordings (0.1% isoflurane) of motor responses (10 trials for each pressure step) expressed as a success percentage; the dashed gray line shows the projection of the motor threshold on the sigmoidal curve; the dashed gray square shows pressures retained for MEP analysis. (C) Electromyographic recordings (1% isoflurane) of motor responses for pressure steps above 80% motor success. (D) Focality of the US beam at peak negative pressure 400 kPa (1 mm at 400 kPa, 2.5 mm from 350 kPa to 400 kPa). EMG: electromyographic, MEP: motor evoked potential. ** p<0.01.

In a first subset of twenty mice processed under 0.1% isoflurane (26), 10 USNS pulses (10-sec apart) were delivered for each acoustic pressure step through a threshold-hunting algorithm (27).

Motor responses in the right forepaw (i.e. contralateral to M1) were assessed without electrodes; overall, the success rate of motor responses increased with pressure, though a total failure of motor responses appeared at 50 kPa (Figure 1B). The motor threshold (MT, i.e. 50% success rate) appeared at 250 kPa. High-success rates (>80%) were found from 350 kPa (81.5±3.9%) to 500 kPa (98.5±0.8%). The experimental data was fit into a sigmoidal curve that followed classical dose-response relationship (26). High-success rates approaching the plateau of the curve (>350 kPa) were retained for further analysis.

The strength and reliability of high-success pressures were evaluated in a second subset of ten distinct mice processed under deeper anesthesia (1% isoflurane) and implanted with subdermal electrodes in the right brachioradialis muscle group (28). Electromyographic (EMG) recordings revealed different strength and reliability of pressures ranging from 350 kPa to 500 kPa. A significant peak average amplitude of motor evoked potentials (MEPs) was obtained at 400 kPa (F(2,18)=13, p=0.0003), above 350 kPa (p=0.0026; Figure 1C). A lower coefficient of variation was also found at 400 kPa (relative standard deviation/RSD=32.4%) in comparison to lower or higher-pressure steps (at 350 kPa, RSD=76.4%; see Table I). The lateral resolution of the US beam at 400 kPa was further estimated between 2.5 and 1 mm, showing greater interest for a focal stimulation of the prefrontal cortex (Figure 1D). Based upon the overall data obtained from M1 stimulation, the optimal US parameters for single-pulse USNS were determined at 400 kPa (160-msec pulse length), displaying criteria of motor success (89±2.7%), strength (MEP=63.7±6.5 µVpp), reliability (RSD=32.4%) and focality (1-2.5 mm) compared to lower or higher-pressure steps.

**Table I.** Determination of optimal US parameters. For all criteria, a peak negative pressure of 400 kPa satisfied high motor success, higher MEP amplitude and lower variability. Statistical differences (post-hoc Tukey) with other pressure steps assessed above MT are shown on the right. MT: motor threshold, PNP: peak negative pressure, RSD: relative standard deviation, RM ANOVA: repeated measures analysis of variance. * p<0.05, ** p<0.01.

|  | PNP (kPa) | motor success (%)<br>mean±SEM | MEP amplitude (μVpp)<br>mean±SEM | Coefficient of variation (RSD) | RM 1-way ANOVA<br>F(2,18)=13<br>p=0.0003 |
| --- | --- | --- | --- | --- | --- |
| below MT | 50 | 0±0 | 14.2±2.67 | 59.57% |  |
|  | 100 | 3.5±2.44 | 15.9±3.02 | 60.01% |  |
|  | 150 | 14.5±5 | 13.4±1.41 | 33.36% |  |
|  | 200 | 36±7.69 | 14.8±1.8 | 38.56% |  |
| MT | 250 | 53±6.77 | 28.2±5.72 | 64.11% |  |
| above MT | 300 | 77±5.03 | 38.2±8.65 | 71.57% | **0.0035** ** |
|  | 350 | 81.5±3.93 | 38.9±9.4 | 76.40% | **0.0026** ** |
|  | **400** | **89±2.7** | **63.8±6.53** | **32.38%** | - |
|  | 450 | 97±1.28 | 41.2±7.93 | 60.83% | **0.013** * |
|  | 500 | 98.5±0.82 | 45.3±9.66 | 67.45% | 0.057 |

### Acute repeated USNS evokes neural activity in the infralimbic cortex

The functional focality and effects of acute repeated USNS was evaluated in eight mice under 1% isoflurane. From single-pulse USNS data and previous studies (16), the pattern of acute USNS was constructed as a 10-min session of 60 USNS pulses (1 pulse: 400 kPa, 160-msec) repeated at a rate of 0.1 Hz and delivered over the prefrontal cortex under 1% isoflurane anesthesia. To target specifically the infralimbic cortex (IL), i.e. the sgACC equivalent in rodents (17), the center of the collimator was positioned at bregma +2 mm on the stereotaxic frame (29). Based on the lateral resolution of the US beam at 400 kPa, the transducer was aimed so to focus the stimulation on the desired structure (Figure 2A). Four mice received acute USNS while four mice received one sham session (“Sham”) that reproduced the exact experimental conditions but with the transducer deactivated.

**Figure 2:**
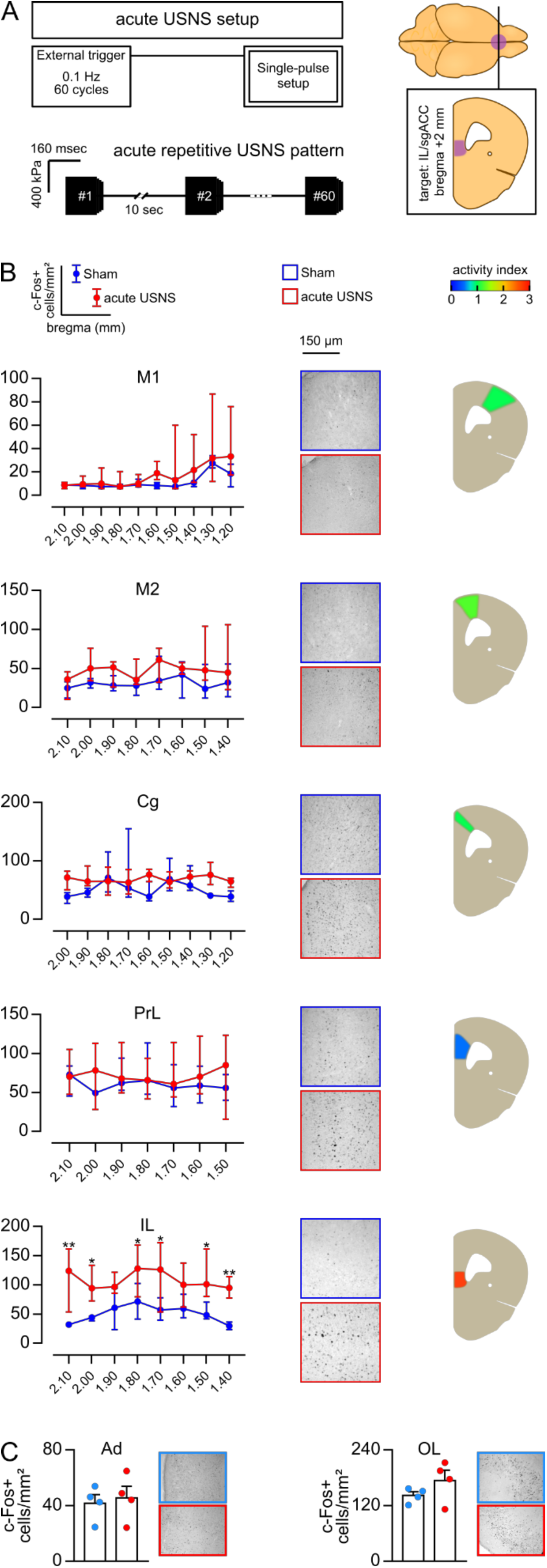
Effects of acute repeated USNS on c-Fos immunolabelling. (A) Schematic view of the system and pattern for acute repeated USNS and brain targeting of infralimbic cortex. (B) (left panel) c-Fos densities (positive cells/mm², median±min/max range) of acute USNS (red) and Sham mice (blue) over the antero-posterior axis of frontal/prefrontal structures expressed in positions relative to bregma (mm); (mid panel) corresponding micrographs (magnificence ×20) of c-Fos cells (black); (right panel) activity index (effect size of acute USNS) showing region-wide increases in c-Fos densities, above 2.5 for the infralimbic cortex. (C) c-Fos densities and corresponding micrographs (×20) for olfactory areas and the auditory cortex (mean±SEM). acute USNS: acute repeated USNS, sgACC: subgenual anterior cingulate cortex, M1: primary motor cortex, M2: secondary motor cortex, Cg: cingulate cortex, PrL: prelimbic cortex, IL: infralimbic cortex, OL: olfactory areas, Ad: auditory cortex. * p<0.05, ** p<0.01.

The immunolabelling of reactive c-Fos neurons revealed that acute USNS elicited neural activation in prefrontal regions. The activity index (i.e. the effect size of acute USNS) was mildly increased in M1 (0.96), motor cortex M2 (1.34), cingulate cortex (Cg, 1.02) and prelimbic cortex (PrL, 0.52), despite showing no statistical difference with Sham mice (Figure 2B). Otherwise, the activity index was further increased in IL (2.67) and multiple analyses revealed that acute USNS elicited significant c-Fos activity along antero-posterior axis of the structure (F(1,3)=14.6, p=0.032). At distance from prefrontal structures, acute USNS did not evoke specific c-Fos activity in the olfactory areas (OL) nor in the auditory cortex (Ad; Figure 2C). In connected subcortical regions, acute USNS induced significant c-Fos activity in subfields of the dorsal/ventral hippocampus (Table II).

**Table II.** c-Fos activity evoked by acute repeated USNS. The differences between means, the student t-tests and the corresponding p-values are presented for each cerebral region compared between one-session USNS and one-session Sham mice. Ad: auditory cortex, AMY: amygdala, BLA: basolateral amygdala, CeA: central amygdala, DG: dentate gyrus, dHPC: dorsal hippocampus, OL: olfactory areas, vHPC: ventral hippocampus. * p<0.05, ** p<0.01.

|  |  | acute USNS vs Sham |  |  |
| --- | --- | --- | --- | --- |
| Area |  | Difference between means | t-test | p-value |
| OL |  | 32±23 | t(6)=1.4 | 0.2149 ns |
| Ad |  | 3.7±10 | t(6)=0.36 | 0.7316 ns |
| AMY | BLA | 9.5±11 | t(6)=0.9 | 0.4012 ns |
|  | CeA | -17±11 | t(6)=1.6 | 0.1646 ns |
| dHPC | DG | 59±34 | t(6)=1.7 | 0.1321 ns |
|  | CA3 | 86±18 | t(6)=4.9 | 0.0026 ** |
|  | CA1 | 37±14 | t(6)=2.5 | 0.0438 * |
| vHPC | DG | 49±13 | t(6)=3.8 | 0.0087 ** |
|  | CA3 | 40±13 | t(6)=3 | 0.0230 * |
|  | CA1 | 46±22 | t(6)=2.1 | 0.0806 ns |

### Chronic repeated USNS treatment alleviates anxiety-related behaviors

The effects of chronic repeated USNS treatment was evaluated in the main cohort of fifty-three mice subjected to the UCMS regimen from day 0 to day 35. At day 7, UCMS mice were significantly different from non-stressed mice in terms of coat state deterioration, a standard measure of UCMS onset and evolution (30) (Supplementary C); from this timepoint, UCMS mice were semi-randomly distributed into the four following treatment/control groups. From day 29 to day 33, chronic USNS (i.e. 5 sessions of acute USNS) was applied to fifteen mice while fourteen mice followed a sham condition (“Sham”). From day 38 to day 43, mice were tested for depressive-like and anxiety-related behaviors in several paradigms. The effects of chronic USNS were assessed against a classic AD compound (fluoxetine, “Flx”), administered chronically through drinking water from day 7 on and controlled (vehicle, “Veh”; Figure 3A). To provide ground values, a group of twelve non-stressed, untreated mice (“Naive”) were processed at the same age through the same behavioral paradigms.

**Figure 3.**
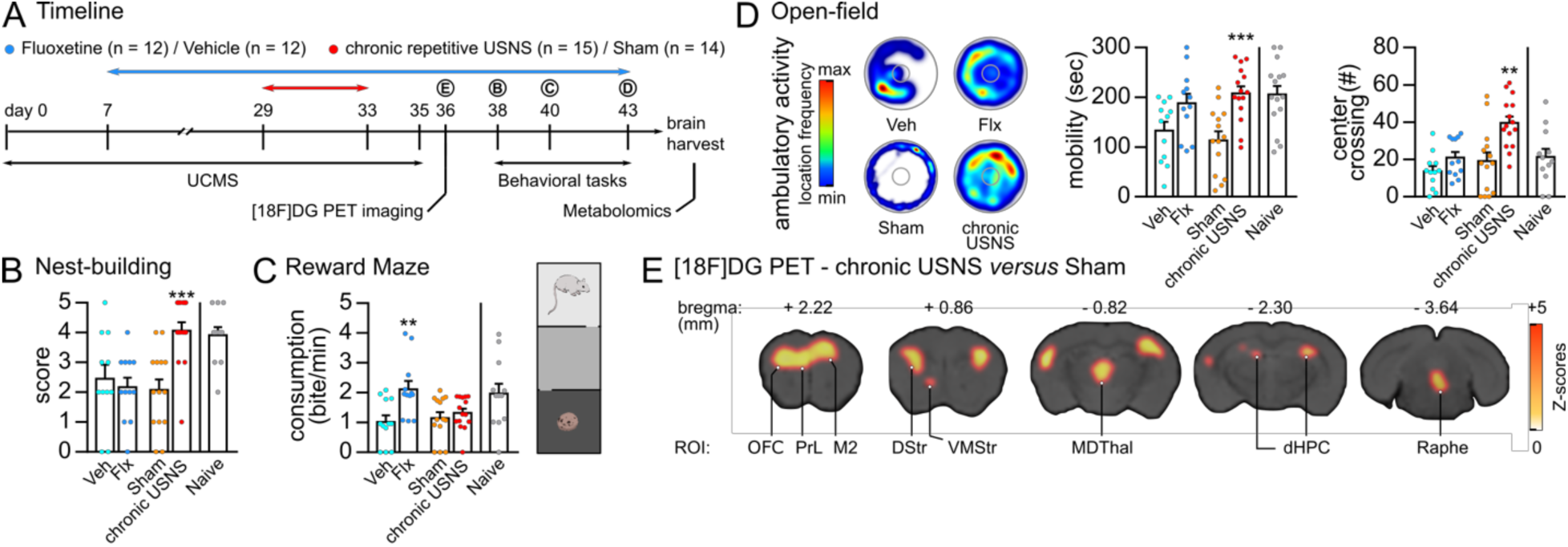
Effects of chronic repeated USNS treatment on behaviors and [^18^F]-FDG uptakes. (A) Experimental timeline for the main cohort (n=53). (B) Nest-building score (scale from 1 to 5). (C) Reward-maze test (5-min duration) apparatus and behavioral measures (consumption of the reward in bites per minute). (D) Open-field task (5-min duration); heatmaps of the cumulative position in the field are extracted from one representative individual of each group; behavioral measures are the mobility (in seconds spent moving) and the occurrence of crossing the 10-mm wide center (count, #). (E) [^18^F]-FDG microPET imaging of chronic USNS versus Sham mice. ROI: regions of interest, chronic USNS: chronic repeated USNS treatment, OFC: orbitofrontal cortex, PrL: prelimbic cortex, M2: secondary motor area, DStr: dorsal striatum, VMStr: ventromedial striatum, MDThal: mediodorsal thalamus, dHPC: dorsal hippocampus, Flx: fluoxetine, Veh: vehicle. Data are expressed as mean±SEM. * p<0.05, ** p<0.01, *** p<0.001.

Overall, both treatments modified behaviors to a different extent. The ability for nest-building, a daily-living measure classically affected by chronic stress (31, 32), was modified by treatments (F (3, 49)=7.9, p=0.0002); chronic USNS mice built the nest faster than Sham mice (p=0.0005), whereas Flx mice did not perform over Veh mice (Figure 3B). Mice were then tested for depressive-like behaviors in a reward-maze paradigm, built of three successive chambers with a palatable biscuit laid in the center of the furthest (30). The relative consumption of the reward, depending on the latency to reach said reward, was modified by treatments (F (3, 49)=5.2, p=0.0032), but only in Flx mice over Veh mice (p=0.042), while chronic USNS induced no behavioral modification. Anxiety-related behaviors were assessed in an open-field task, where both treatments enhanced qualitatively the ambulatory activity in the field (Figure 3D, heatmaps). Mobility in the open-field was significantly increased in chronic USNS mice (F (3,49)=7, p=0.0005) over Sham mice (p=0.014), whereas Flx mice did not statistically differ from Veh mice despite a qualitative increase of mobility. Moreover, chronic USNS increased the number of center-crossings (F (3, 49)=10, p<0.0001) over Sham mice (p=0.0008), while Flx did not significantly affect the measure (Figure 3D).

### Chronic repeated USNS treatment modifies short-term cortical and subcortical brain metabolism

Seventy-two hours past the last chronic USNS session, changes in brain metabolic activity were assessed using [^18^F]-FDG microPET imaging in awake animals. Compared to Sham mice (n=10/group), USNS revealed significantly increased metabolic activity in frontal cortical regions including the prelimbic/M2 area, but also in the orbitofrontal regions. In addition, chronic USNS increased significantly the metabolic activity in distant subcortical regions such as the dorsal and ventral striata, the thalamus, the dorsal part of the hippocampus, the periaqueductal gray matter (PAG) and the raphe nuclei (Figure 3E, Table III). Other regions underneath the target (e.g., anterior olfactory regions and dorsal peduncular cortex), did not display modified metabolic activities, neither did laterally adjacent regions such as somatosensory cortices, or the direct antero-posterior vicinity.

**Table III.**
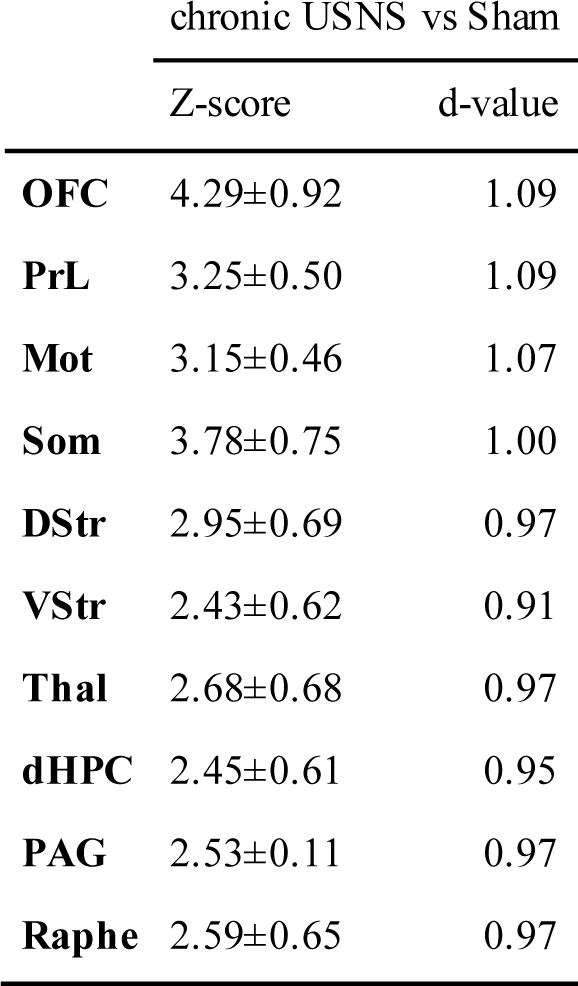
Statistical significances for inter-group comparisons in [^18^F]DG uptake. The Z-scores and d-values are presented for each significant difference observed between chronic USNS and Sham mice. Large effect sizes correspond to d-values comprised between 0.80 and 1.20 (61). dHPC: dorsal hippocampus, DStr: dorsal striatum, Mot: motor cortex, OFC: orbitofrontal cortex, PAG: periacqueductal gray, PrL: prelimbic cortex, Raphe, Som: somatosensory cortex, Thal: thalamus, VStr: ventral striatum.

| chronic USNS vs Sham |  |  |
| --- | --- | --- |
|  | Z-score | d-value |
| **OFC** | 4.29±0.92 | 1.09 |
| **PrL** | 3.25±0.50 | 1.09 |
| **Mot** | 3.15±0.46 | 1.07 |
| **Som** | 3.78±0.75 | 1.00 |
| **DStr** | 2.95±0.69 | 0.97 |
| **VStr** | 2.43±0.62 | 0.91 |
| **Thal** | 2.68±0.68 | 0.97 |
| **dHPC** | 2.45±0.61 | 0.95 |
| **PAG** | 2.53±0.11 | 0.97 |
| **Raphe** | 2.59±0.65 | 0.97 |

### Chronic repeated USNS treatment modifies long-term metabolomics in cortical and subcortical regions

Ten days pas the last chronic USNS session, significant modifications were found when compared to Sham mice (n=8/group) in the metabolome of interconnected brain regions involved in the UCMS model (23) and MD (6): the Cg, the prelimbic/infralimbic cortex (PrL/IL), the amygdala and the hippocampus.

In cortical regions, chronic USNS had a relatively low impact on the metabolome of the Cg, since no metabolic pathway was found significantly disturbed, despite 9 metabolites showing significantly different concentrations compared to Sham mice (Table IV). In the PrL/IL of chronic USNS mice, 6 metabolites were significantly modified, including the decrease of glutamic acid/glutamate concentrations (Table IV). Five metabolic pathways were found disrupted, including arginine-proline metabolism (false discovery rate (FDR)=7.8×10^−4^), alanine metabolism, aspartate and glutamate (FDR=0.016), glutathione metabolism (FDR=0.017), histidine metabolism (FDR=0.024), glycine metabolism, serine and threonine (FDR=0.035) and aminoacyl-tRNA biosynthesis (FDR=0.038; Table V).

**Table IV.** Metabolites significantly modified by USNS in the Cg and the PrL/IL. FC: fold change. The total number of modified metabolites is given for both regions.

| Cingulate cortex |  |  | Prelimbic/Infralimbic cortex |  |  |
| --- | --- | --- | --- | --- | --- |
| Metabolites (n=86) | FC | p-value | Metabolites (n=83) | FC | p-value |
| **Phenylacetic acid** | 2.50 | 0.002 | **Trans-aconitate** | 1.37 | 0.009 |
| **Glucosamine 6-phosphate** | 2.60 | 0.014 | **4-Hydroxy-D-proline** | 2.88 | 0.020 |
| **Lipoamide** | 1.95 | 0.029 | **Homovanillate** | 0.52 | 0.020 |
| **Pyridoxamine** | 2.29 | 0.040 | **Glutamic acid** | 0.37 | 0.029 |
| **Cysteine** | 0.47 | 0.040 | **Hypoxanthine** | 2.79 | 0.040 |
| **Guanosine 5'-monophosphate** | 2.13 | 0.040 | **1-NH2-Cyclopropane-1-carboxylate** | 1.48 | 0.040 |
| **Glutamine** | 2.12 | 0.040 |  |  |  |
| **Homocystine** | 0.77 | 0.040 |  |  |  |
| **Creatine** | 1.16 | 0.040 |  |  |  |

chronic USNS vs Sham

**Table V.**
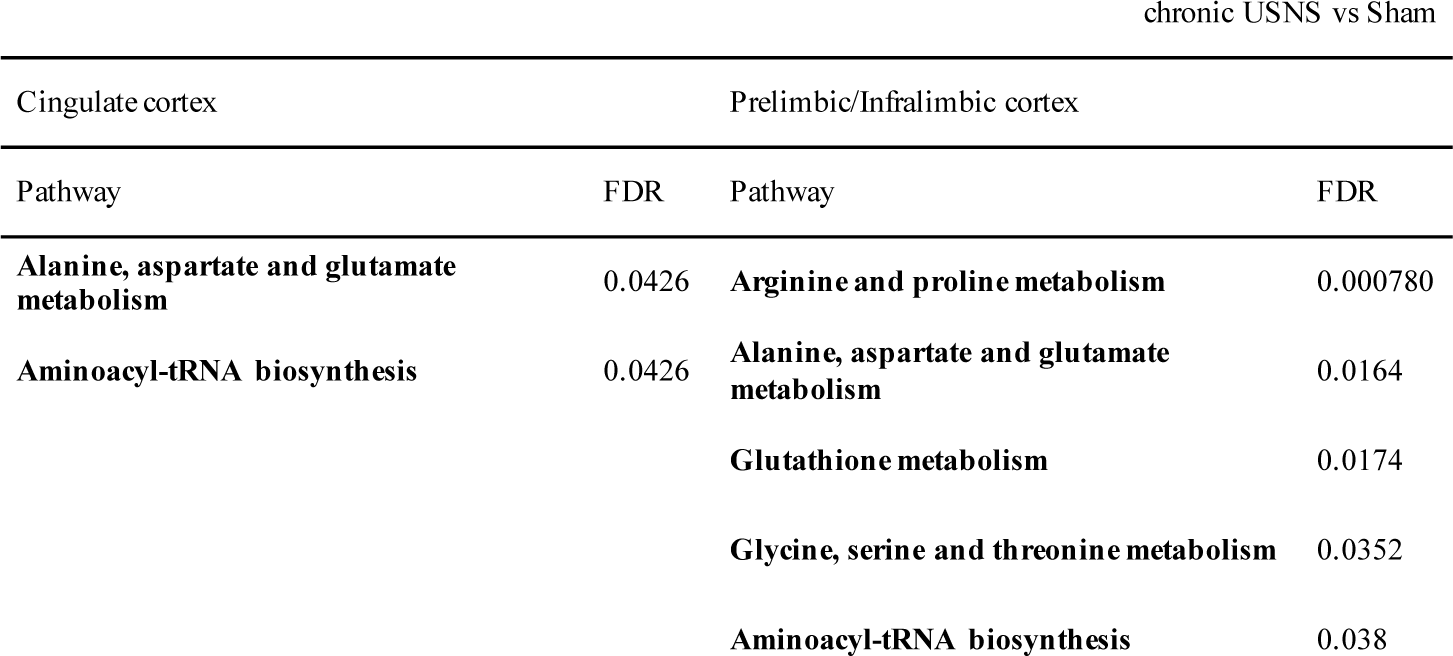
Metabolic pathways significantly modified by USNS in the Cg and the PrL/IL. FDR: false discovery rate.

| Cingulate cortex |  | Prelimbic/Infralimbic cortex |  |
| --- | --- | --- | --- |
| Pathway | FDR | Pathway | FDR |
| **Alanine, aspartate and glutamate metabolism** | 0.0426 | **Arginine and proline metabolism** | 0.000780 |
| **Aminoacyl-tRNA biosynthesis** | 0.0426 | **Alanine, aspartate and glutamate metabolism** | 0.0164 |
|  |  | **Glutathione metabolism** | 0.0174 |
|  |  | **Glycine, serine and threonine metabolism** | 0.0352 |
|  |  | **Aminoacyl-tRNA biosynthesis** | 0.038 |

In subcortical regions, the amygdala was impacted where 7 metabolites were found significantly modified (Table VI). Nine metabolic pathways were found significantly disturbed including aminoacyl-tRNA biosynthesis (FDR=4.48×10^−5^), abnormalities in lysine biosynthesis (FDR=0.046), in addition alanine, aspartate and glutamate metabolism (FDR=0.011); Table VII). In the hippocampus, 5 metabolites were found significantly modified (Table VI). Five metabolic pathways were also significantly disrupted including aminoacyl-tRNA biosynthesis (FDR=2.16×10^−7^), alanine-aspartate-glutamate and nitrogen metabolisms (FDR=0.004), arginine-proline metabolisms (FDR=0.013), and glutathione metabolism (FDR=0.023; Table VII). Despite long-term changes in the metabolome of the hippocampus, no effects of chronic USNS were found on the proliferation of newborn neurons in the dentate gyrus of the hippocampus (DG), a key mechanism to the function of classic ADs (Supplementary B).

**Table VI.**
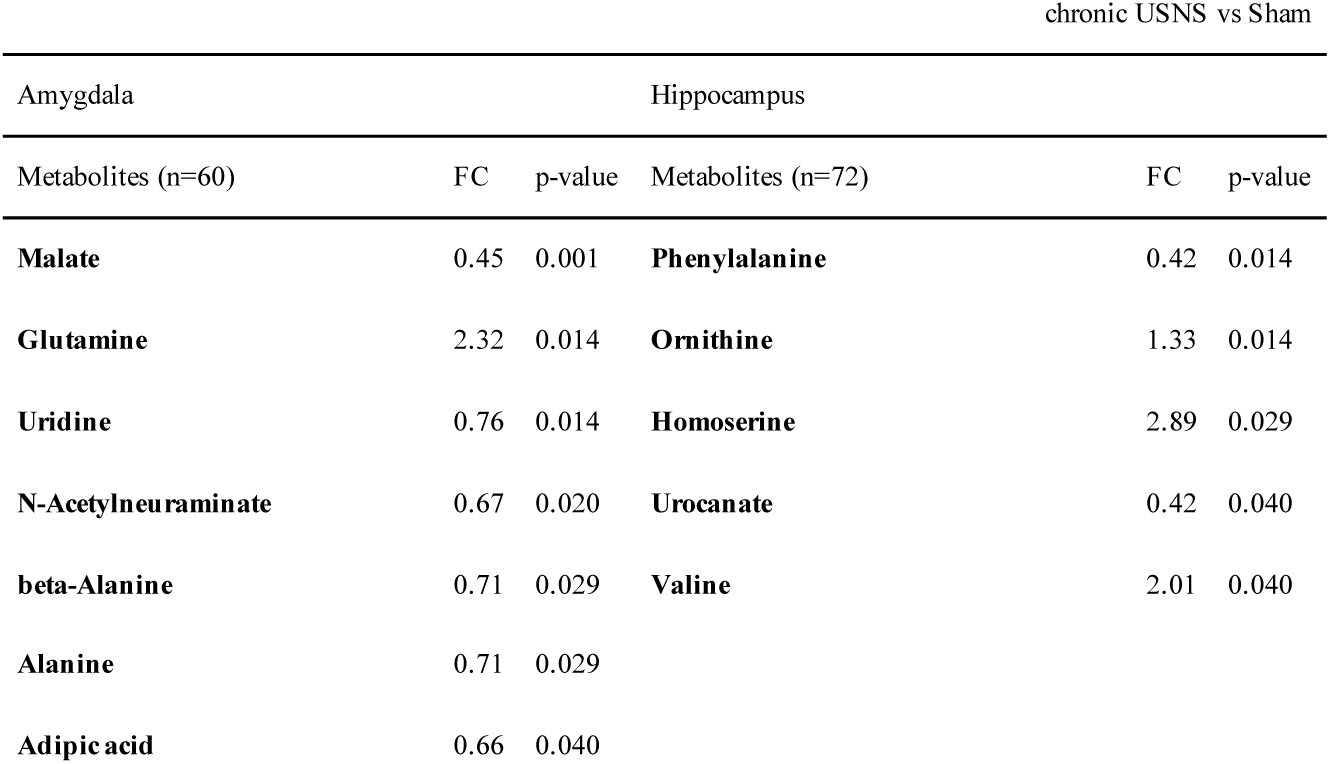
Metabolites significantly modified by USNS in the amygdala and the hippocampus. FC: fold change. The total number of modified metabolites is given for both regions.

**Table VII.** Metabolic pathways significantly modified by USNS in the amygdala and the hippocampus. FDR: false discovery rate.

| Amygdala |  | Hippocampus |  |
| --- | --- | --- | --- |
| Pathway | FDR | Pathway | FDR |
| Aminoacyl-tRNA biosynthesis | $4.48 \times 10^{-5}$ | Aminoacyl-tRNA biosynthesis | $2.16 \times 10^{-7}$ |
| Arginine and proline metabolism | 0.000183 | Alanine, aspartate and glutamate metabolism | 0.00416 |
| Nitrogen metabolism | 0.00246 | Nitrogen metabolism | 0.00416 |
| beta-Alanine metabolism | 0.00246 | Arginine and proline metabolism | 0.0128 |
| Alanine, aspartate and glutamate metabolism | 0.0114 | Glutathione metabolism | 0.0229 |
| Histidine metabolism | 0.0121 |  |  |
| Glutathione metabolism | 0.0121 |  |  |
| Glycine, serine and threonine metabolism | 0.024 |  |  |
| Ubiquinone and other terpenoid-quinone biosynthesis | 0.0264 |  |  |
| Lysine biosynthesis | 0.0466 |  |  |

## Discussion

Numerous brain structures are known actors in the pathophysiology of MD, yet the growing need to act therapeutically on these regions remains only partially answered by current ADs and neurostimulation techniques. The present study used repeated mechanical US waves to non-invasively target the IL/sgACC in a mouse model of MD. Chronic repeated USNS treatment impacted various behavioral endpoints induced by the UCMS regimen, which was associated to short-term changes in brain metabolic activity on the site of stimulation (prefrontal cortex and its close surroundings) but also at distant, connected limbic regions, such as the striatum, the dorsal hippocampus and the raphe nucleus. Measures of well-being (nest-building) and anxiety-related behaviors (open-field task) were ameliorated by chronic USNS, while items relative to anhedonia and reward-seeking were not readily modified compared to classic SSRI treatment (fluoxetine). Metabolites were modified at long-term in cortical regions on target site and in subcortical regions in the amygdala and the hippocampus, involving glutamate pathways that might correlate to longer-term changes in brain plasticity.

The ability of US waves to reliably stimulate cortical regions through the cranium has been reproduced in the current study with single-pulse USNS of the primary motor cortex M1. While the current data reproduced previous findings in terms of the sigmoidal relationship of the pressure to the motor success at low dose anesthesia (26), the quantitative analysis of MEPs showed that the strength of the stimulation and its reliability over multiple stimulations might depend on other factors than acoustic pressure alone. Consistent rising of motor success with acoustic pressure points toward an activation of glutamatergic neurons, but the inconsistency of MEP amplitudes with higher pressures (>400 kPa) could reflect the recruitment of different neural populations depending on US parameters, even though desensitization biases were controlled with a 10-sec spacing of US pulses (16). Such variations upon intensity/pulse length were observed for different neurostimulation approaches (i.e. rTMS) and might apply to USNS (33, 34). Thus, the determination of optimal US parameters in terms of pressure/pulse length appears pivotal to induce reliable brain stimulation with US waves. Furthermore, the deeper anesthesia used during the MEP procedure (1% isoflurane) could have modified the quality of motor signals at pressures above 400 kPa and so despite higher motor success rates at 0.1% isoflurane; the concentrations used in most studies for MEP analysis (<0.25 %) were too low to fit those of the chronic USNS paradigm (1%), where a semi-awoken state could impart anxiety biases. We thus reckoned that the current findings at 1% isoflurane could be more readily transposed to the treatment condition of chronic USNS. Concurrently, motor responses could be diminished by slow-moving cortical spreading depolarization (14), occurring for lastly tested pressures (500 kPa), which further underpin the need to finely tune US pulses in terms of intrinsic parameters and sustained effects (35).

When applied for one unique session, acute USNS was able to evoke neural activity in the IL of mice with little spatial inaccuracy at stimulation site: surrounding brain regions such as the PrL or the Cg, M2 and M1 were not readily affected by the stimulation in comparison to sham-treated mice. On the other hand, distant hippocampus regions were affected by acute USNS. Given the geometric properties of the US beam, the results suggest that, similarly to standard M1 stimulation, a threshold pressure appears for patterns of repeated stimuli, under which the activation of a brain region was not seen on immediate c-Fos labelling; the lack of c-Fos activation in other prefrontal regions suggests that the spatial resolution of acute USNS is not directly correlated to its functional resolution. The narrower range of effective acoustic pressures could be responsible for the specific stimulation of the IL (1-mm wide at 400 kPa). Furthermore, the auditory cortex was not affected by acute USNS, which further supports a functional targeting of the IL (36, 37). Because c-Fos labelling does not discriminate between glutamatergic and GABAergic neurons, the results show that brain regions might react differently to acute USNS. As pulsed neurostimulation is described to preferentially act on axons rather than somas (38), the reactivity of a brain region to the application of USNS might depend on the orientation of the stimulation and the frequency of pulse repetition as it has been shown for rTMS (39), highlighting the discrepancy between geometric and functional focality when applying repeated, sustained stimulation.

When applied chronically for 5 days, behavioral measures were modified by repeated USNS. As an indicator of well-being in rodents, the onset of nest-building was reduced by chronic USNS, while it was not affected by fluoxetine (similar effects of the drugs were previously reported (31)). Because such daily-living activities can be negatively regulated by chronic stress and MD, chronic USNS showed beneficial effects on this aspect of the UCMS model. Furthermore, chronic USNS reduced anxiety-like behaviors: measures of mobility and center-crossings were found significantly different from sham-treated mice in the open-field task. On the other hand, the anhedonic measure of reward consumption was not ameliorated and did not match the results of fluoxetine-treated mice. It was reported that a single session of isoflurane anesthesia could induce antidepressant-like effects and increase glutamatergic transmission in the hippocampus (40), however, the sham-treated group displayed baseline behaviors statistically equal to vehicle-treated mice, suggesting that the present results are specific to the chronic USNS condition. As opposed to fluoxetine, where a variability of anxiolytic effects is reported (5), chronic USNS might act differently from classic SSRIs, involving top-down mechanisms of the prefrontal cortex over subcortical regions. The high comorbidity of MD with anxiety disorders, further disabling for patients, is not fully treated by classic SSRIs (5), underpinning different therapeutic paths for both anxiety and depressive-related symptoms. Classic AD compounds also show different effects based upon the duration of administration, with a lower impact at an acute/sub-chronic stage of medication (41). Furthermore, the current results show different levels of response to fluoxetine, with bimodal distributions appearing in behavioral measures; this might reflect similar resistance mechanisms observed in patients with MD, or the inadequacy of fluoxetine for some subjects. On the other hand, chronic USNS mice displayed lower spread distributions, which could suggest that the technique imparts beneficial effects independently from inter-individual variability and in a shorter timespan following treatment. Finally, because anxiety-related and anhedonic features do not rely on the same brain correlates, the anxiolytic effects of chronic USNS might appear early following treatment. However, the antidepressant effects, although only slightly impacted, could develop after a longer period and depending on long-term changes in metabolic activity.

The direct aftermath of chronic USNS was evaluated with microPET imaging 72 hours past the last treatment session. The uptake of [^18^F]-FDG was increased near the target site at bregma +2 mm in prefrontal regions (mostly the PrL and the orbitofrontal cortex (OFC)) and in M2. Because this latter region cannot presently be linked to brain modifications induced by the UCMS regimen and more broadly in MD, the increased activity might be a direct effect of chronic USNS, either showing spatial biases or a different functional focality when repeated chronically as opposed to acute USNS. Other regions within a 1.5-mm radius of the stimulation site did not appear modified during microPET imaging, suggesting that presumably only the most intense portion of the US beam acts upon the target also in this timespan. The brain metabolic activity was also increased in distant regions (striatum, thalamus, dorsal hippocampus, raphe nucleus and PAG), which could mean that chronic USNS acted at distance from the prefrontal target through functional connectivity, which is supported by the effects of acute USNS on the hippocampus. The effective stimulation of such projections might take part in the behavioral effects of chronic USNS: hippocampal-prefrontal communication is considered crucial in the pathophysiology of MD (42), and the connectivity of the PrL/IL cortex to the raphe nucleus has been identified in rats to play a role in behavioral control of stressors (43), which could relate to the decrease of anxiety-related behaviors. Furthermore, PET-Scan imaging revealed that MD was associated with an hypoactivation of frontal regions (44) and that therapeutic response to fluoxetine could revert that type of metabolic changes by increasing cortical activity (45); in the same study, failure to respond to classic SSRI treatment was associated with an absence of cortical modification. The effects of USNS on cortical metabolism, and more specifically the prefrontal regions, could have participated in the modification of the UCMS-induced phenotype. Cortical changes could be at play in the top-down regulation of anxiety-related behaviors but would require larger sample size to argue.

Ten days past the last treatment session, cortical (Cg, PrL/IL) and subcortical structures (amygdala and hippocampus) displayed significant changes in metabolic pathways following chronic USNS. Likewise, glutamate pathways were similarly modified in all studied regions, and more specifically, decreased levels of glutamate were observed in the PrL/IL. In previous studies, glutamate levels were reported to increase in the serum and in frontal regions of MD patients (46, 47). In rodents, glutamate is implicated in the expression of depressive-like and anxiety-related behaviors (48). Furthermore, glutamate pathways in the hippocampus, but not the prefrontal cortex, might be pivotal to antidepressant response in SSRI-treated mice (49). The current findings thus show that the effects of chronic USNS on glutamate pathways could be associated to a therapeutic response similarly to classic fluoxetine treatment, although both treatments did not modify the same behaviors in the UCMS model. In addition, our results showed that other hippocampal functions, such as neurogenesis, were not modified by chronic USNS, which further supports that the behavioral effects were mainly observed on anxiety and not directly on anhedonic features. In the present study, few metabolic modifications were found in the Cg of stimulated mice, even though this structure was in the propagation path of the US beam during stimulation. Direct interneuron regulation between the structures could have been at play and thus participated in reversing prefrontal abnormalities induced by UCMS (20). Glutamate variations in other brain regions, such as the PAG, have also been linked to depressive-like behaviors and chronic stress (50). Analysis of this region was beyond the scope of our study, although [^18^F]-FDG metabolism in the PAG was modified by chronic USNS treatment.

We should emphasize that microPET imaging was performed at day 36 while behavioral tests were carried out from day 38 to day 43, and metabolomic analyses performed after brain harvest at day 43. Further investigation of the effects of chronic USNS treatment on the brain is required to explore potential effects and therapeutic outcomes. Stimulation of the M1 highlighted the need to identify a precise threshold that does not exceed or fall short the efficient intensity. The metabolomic analyses suggested that USNS did not induce excitotoxic effects on the target as common inflammatory metabolites were not observed in screened regions (51, 52). Because the technique does not rely on electromagnetic waves, US transducers could be associated with calcium imaging without interference and adapted for fMRI to further the comprehension of its mechanisms. On-line functional imaging of the brain tissue reaction to USNS could reveal if specific neural populations are modulated (e.g. pyramidal/interneurons) as a function of stimulation duration and intensity (34). Future research in US transmission through the human skull could bring forward new findings on the effects of USNS in psychiatric disorders and its potential use in regulating impaired brain networks.

## Conclusions

Major depression is one of the main factors contributing to the Global Burden of Disease. Current treatment strategies of major depression have shown limitations, such as inaccuracy and invasiveness. In this study, we evaluated the potential of US neurostimulation (USNS) in an unpredictable chronic mild stress (UCMS) model. In comparison to pharmacological treatment (fluoxetine), the results showed that selected US application on the prefrontal cortex counteracts behavioral modifications induced by the UCMS regimen and decreases anxiety-related behaviors to a greater extent than classic fluoxetine treatment. Next to these effects, chronic repeated USNS treatment triggered the activation of various brain regions including regions at distance from the targeted zone as confirmed by microPET imaging and metabolomic analyses. These results demonstrate the potential of USNS as a therapeutic tool for major depression.

## Materials and Methods

### Experimental layout

Eighty-three male BALB/cByJRj mice were obtained from Janvier Labs (Le Genest-Saint-Isle, France), aged 9 weeks (29±1.5 g) at the beginning of the experiments. Animals were housed in standard condition (12:12 light-dark cycle, room temperature 22±2° C, free access to food and water). Thirty mice were used for setting-up optimal US neurostimulation (USNS) parameters with motor response measurements, determined with: 1) electrode-free video recordings of the targeted limb (n=20) and 2) electromyography (n=10). The remaining cohort (n=53) was divided in four distinct treatment groups (Veh, Flx, Sham, chronic USNS) that underwent 35 days of the unpredictable chronic mild stress regimen (UCMS) to induce depressive-like behaviors. Then, from day 7 on, mice from the Veh (vehicle) and the Flx (fluoxetine) groups were chronically treated through drinking water respectively with water alone (n=12) or 15 mg/kg fluoxetine (n=12). Every day from day 29 to day 33, mice from the chronic USNS and the Sham groups were either treated with repeated USNS under 1% isoflurane gaseous anesthesia (n=15) or solely anesthetized (n=14). The whole cohort underwent behavioral tasks from day 38 to day 43 to test for depressive-like and anxiety-related behaviors. To assess the underlying mechanisms, a subset of mice (n=10/group) of the Sham and chronic USNS groups were further analyzed: at day 36 (72 hours past the last session of chronic USNS/Sham), chronic USNS mice were scanned with [^18^F]-FDG microPET imaging and compared to Sham mice. At day 43 (10 days past the last session of chronic USNS/Sham), brains were harvested to observe metabolome changes that occurred in chronic USNS (n=8/group) versus Sham mice (n=8/group). Animals that went for metabolomics were also scanned with microPET imaging. All experiments were compliant with Directive 2010/63/EU guidelines on animal ethics.

### Brain stimulation setup

Ultrasound stimuli were generated using a single-element transducer focused at 65 mm (active diameter of 38 mm, Imasonic, Besançon, France) with a central frequency of 500 kHz and a fractional bandwidth of 58%. The generated acoustic pressures were measured in a degassed water tank using a calibrated hydrophone (HGL 200, ONDA, Sunnyvale, CA, USA) positioned at the focus. The attenuation coefficient of mice skulls (n=7) was estimated ex-vivo using standard through-transmission insertion loss techniques (53). Briefly, the skull was placed along the ultrasound beam between the source transducer and the hydrophone. At 500 kHz, the attenuation coefficient was 6.32±2.18% (mean±SD). This coefficient was used to assess the derated acoustic pressure inside the brain during US stimulation assuming the attenuation in the mouse brain tissue is negligible.

The transducer was positioned on the mouse brain with an appended plastic column, filled with degassed water. A distal collimator of 10 mm was sealed with polyethylene and coupled with centrifuged US gel to the shaved cranium of the animal. Throughout all US-related procedures, mice were anesthetized with 1.8 liters per minute gaseous isoflurane (2.5% induction, 1% maintenance or 0.1% maintenance for electrode-free M1 stimulation; halogenated ether, Aerrane, Baxter SAS), placed in a stereotaxic frame (SM-6M-HT, Narishige), the head fixed with auxiliary ear bars (EB-5N, Narishige), while the transducer column was operated by a custom-made 3-axis stereotaxic manipulator (27). Electrical signal was generated from a function generator (Agilent, Santa Clara, CA, USA) and then amplified using a power amplifier (500W ADECE, Artannes sur Indre, France; Figure 1A).

### EMG recordings

For harvesting motor evoked potentials (MEPs), subdermal electrodes were positioned in the right brachioradialis muscle group (active) and between the 3^rd^ and 4^th^ carpometacarpal joints (27). Signals were acquired, with a sample rate of 2 kHz (PowerLab, AdInstrument, Australia) and analyzed in post-treatment (Labchart 7, AdInstrument, Australia). A band pass digital filter, between 300 Hz and 1 kHz, was applied, then the absolute value was taken on 301 samples. A single-event MEP was monitored up to a 100-msec bin for late polysynaptic waves. Averaged MEP amplitudes were then obtained for each individual regardless of motor success.

### c-Fos analysis of acute repeated USNS

Ninety minutes after a unique session of repeated USNS (“acute USNS”) or only anesthesia (“Sham”, 1% isoflurane), mice were injected (i.p.) with an overdose of pentobarbital (Dolethal), transcardially perfused with 40 mL of saline (0.9% NaCl) to remove the blood reservoir, then perfused with 100 mL of 4% paraformaldehyde (PFA). The brains were harvested, left in 4% PFA overnight then put into a sucrose solution (20%) at 4° Celsius for 48 hours. The brains were then snap-frozen in dry-ice-cooled isopentane and stored at −80° Celsius before the rest of the procedure. To be processed for immunohistochemistry (IHC), the brains were cut into 40-µm coronal sections with a cooled microtome (−20° Celsius, Leica CM 3050 S). A classical method was employed for free-floating IHC. After endogenous peroxydase blockade (20 min, 50% EtOH, 1% H2O2), sections were processed with primary antibodies directed against c-Fos (1:1000, SC-52-G goat polyclonal IgG, Santa Cruz Biotechnologies) in phosphate buffer (PB) 0.1 M, 2% Normal Donkey Serum and 0.1% Triton for 48 hours at 4° C. Then a secondary incubation (1:500, Biotin-SP-conjugated AffiniPure donkey anti-goat IgG, Jackson ImmunoResearch) was performed 2 hours at room temperature. Finally, a standard protocol was used with 1-hour 1%-avidin/1%-biotin complex (Vectastain Elite ABC kit) and 3,3’-di-amino-benzidine revelation (SIGMAFAST™ DAB tablets, Sigma-Aldrich) for 3 minutes.

The immunolabelled sections were observed under a Zeiss Z.2 Imager microscope in transmitted-light mode. Micrographs (magnificence ×10) were exported to the processor ImageJ (53) in grayscale 8-bit format and converted to a binary mask at 60% of background’s mean gray value, which produced a count of c-Fos-positive (c-Fos+) cells in a selected region of interest. To analyze c-Fos activity patterns associated to a USNS or Sham, ubiquitous sections were picked for the olfactory areas (OL), which are paramount to behaviors in rodents, the auditory cortex (Ad) (36, 37), and frontal/prefrontal structures that lay under the US beam or its direct vicinity: the prelimbic cortex (PrL), the infralimbic cortex (IL), the anterior cingulate cortex (Cg), the primary motor cortex (M1) and the secondary motor cortex (M2). Furthermore, subcortical regions connected to prefrontal regions were evaluated: 1) the amygdala (AMY) and its subfields: the basolateral amygdala (BLA) and central amygdala (CeA), and 2) the dorsal/ventral hippocampus (dHipp/vHipp) and its subfields: the dentate gyrus (DG), the CA3 and the CA1. The number of c- Fos+ cells in each region of interest was expressed as normalized cellular densities (c-Fos+ cells/mm²) (Supplementary A).

### Unpredictable chronic mild stress

The main cohort of fifty-three mice followed the UCMS regimen from day 0 to day 35 (Figure 3A), as described previously (54). Briefly, mice were isolated in 24×11×12 cm cages without environmental enrichment and submitted to daily random socio-environmental stressors including exposition to another mouse bedding, removal of sawdust, contention and light-dark cycle perturbations.

### Treatments

Pharmacological treatments (15 mg/kg fluoxetine) were given from day 7 on to achieve chronic administration at week 5 during analyses. Short delays were described before (30) and allow to observe positive effects of the drug in the experimental timespan. At week 1 of UCMS, the coat state, a standard measure of general deterioration due to UCMS, was significantly deteriorated compared to non-stressed mice and UCMS mice were semi-randomly divided into the four treatment groups (Veh, Flx, Sham, chronic USNS; Supplementary C). The chronic repeated USNS treatment was given during the fifth week of the UCMS (stressors: 08:00 – 12:00 AM, treatment: starting at 02:00 PM). To perform chronic USNS, another function generator (Agilent, City, CA, USA), called external trigger (Figure 2A), was used to control the repetition rate of the US stimulus defined in the motor cortex stimulation procedure. A 5 Vpp square wave, set to 0.1 Hz, was used to trigger the main function generator, set for 80,000 cycles (160-msec pulse) of a sine wave (500 kHz). The pattern was repeated for 10 min (60 US stimuli) every 10 sec (0.1 Hz); higher repetition frequencies were reported to blunt cortical response (16). This constituted one treatment session of chronic USNS (or “acute USNS”); one session was applied every day for 5 consecutive days from day 29 to day 33 to constitute the chronic protocol of chronic USNS (total of 300 US stimuli over 60 min). Through stereotaxic framing, the collimator was positioned along the interhemispheric line (midline) at bregma +2 mm, a region that comprises the infralimbic cortex (IL), the rodent’s equivalent of the sgACC connectivity-wise (17).

### Behavioral Measures

Behaviors were assessed during the dark phase of the light-dark cycle, i.e. the active, awoken phase for the animals. The effects of USNS were assessed against those of the fluoxetine treatment from day 38 to day 43 with 1) the nest-building test (55), 2) the reward-maze test (30) and 3) the open-field task.

1. For the nest-building task, mice were moved on day 37 to Makrolon type III cages, then at 07:30 am of day 38, a square pressed cotton (5 cm²) was placed on the sawdust. After 5 hours in standard condition, the state of the nest, built in cotton, was scored on a predefined scale (55). The mice were then put back in their smaller home cage.
2. To test for anhedonic traits, mice were subjected to a reward-maze test (30). The palatable value of the reward was induced by giving a sample every day to the animals from week 2 to week 3. To minimize environmental neophobia, mice were habituated three times to the paradigm on week 4 and the test was done under red light. The apparatus was made of three consecutive chambers (20×20×20 cm) growing darker in color (light gray to black). Common food pellets were removed from the cage lid 1 hour before the test. The mouse was placed in the light chamber and the reward at the center of the black one. The consumption of the reward was measured for up to 5 min and put in relation to the latency of reaching the reward, i.e. the consumption is expressed as the ratio of bites taken into the reward over the time spent with the reward (300 sec minus the latency to reach the reward).
3. To test for anxiety-related behaviors, mice were subjected to the open-field task on day 43. Mice were placed in a brightly lit (200 lux) circular 33-cm wide open-field. The mobility (i.e. the time spent moving in the field, expressed in seconds), the number of center-crossings (center area, d=10 cm) and ambulatory behaviors (expressed in heatmaps as a cumulated path) were measured for up to 5 min. For the latter, the movements of the mice were recorded and tracked with the software EthoVision XT (Noldus Information Technology, Netherlands) to generate individual heatmaps, color-coded for position frequencies over the duration of the test.

### Brain Imaging

At day 36, awake mice were injected with [^18^F]-FDG (18.5 MBq/100g i.p.; Cyclopharma, Tours, France), and placed in their home cage for 45 min. Then, animals were anesthetized using isoflurane 4% (Baxter, Maurepas, France), placed on a heating pad (Minerve, Esternay, France) and centered in the field of view of the Explore VISTA-CT microPET camera (GE Healthcare, Velizy, France). A CT-scan was performed for attenuation correction of PET images and a list-mode PET acquisition of 30 min started 60 min after [^18^F]-FDG injection. After data reconstruction using a 2-D OSEM algorithm, all images were co-registered and normalized for tissue activity in the whole brain. Quantitative results were expressed as mean±SD and were presented on Z-score maps using an array of regions of interest already defined in PMOD v3.2 software (PMOD Technologies Ltd, Switzerland).

During the experiments, the respiratory rate and body temperature of each animal were monitored and kept as constant as possible (70 respirations per minutes and 37°C, respectively). List-mode scans were rebinned into 6 frames of 300 sec, corrected for random, scatter and attenuation, and images were reconstructed using a 2-D OSEM algorithm (GE Healthcare, Velizy, France) into voxels of 0.3875×0.3875×0.775 mm^3^. Data summed over the entire acquisition were used for image registration. Since brain anatomy is very similar for mice of similar weight (56), registration was accomplished as a rigid body transformation, with no warping or scaling. Each summed scan was individually smoothed with a Gaussian filter to improve the signal-to-noise ratio and to reduce the bias of misregistration into template space. For this smoothing, a kernel of 0.6×0.6×0.6 mm^3^ FWHM was used. Each scan was coregistered using PMOD v3.2 software (PMOD Technologies Ltd, Switzerland) to a [^18^F]-FDG PET template in Paxinos coordinates (57) using a mutual information similarity function with Powell’s convergence optimization method (58). The results were visually checked for misregistration. Each summed image was also used for statistical analysis. The regions of interest (ROI) atlas of Mirrione in Paxinos coordinates were merged to create a whole brain mask (WBM). To normalize the [^18^F]-FDG uptake, tissue activity was divided by the whole brain activity, calculated as the average activity in the WBM. Prior to statistical analysis, the WBM was applied over all PET scans to exclude extracerebral regions. The signals extracted using the ROIs on the Z-score maps were considered for further analysis when representing at least 50 contiguous voxels for a statistical threshold set at p<0.05.

### Metabolomics

Each sample was first lyophilized and then weighed precisely in order to finally normalize the results to the dry mass of tissue. Metabolites were then extracted from approximately 1-3 mg of tissue by two successive extractions, after homogenization, with a mixture of methanol/water (1/1, 0.75 mL). After centrifugation, the supernatant was collected, and the solvent evaporated by means of a speedvac. The dry residues were finally taken up in 150 µL of MeOH/H2O (1/1). 10 µL extracts of each sample were pooled to obtain a mixture used as a quality control. Finally, 20 µL were used for LC-HRMS analysis. Fifteen quality control (QC) samples were injected to equilibrate the chromatographic system before each analyses batch. The running order of samples was randomized, and QCs were analyzed every 10 samples. The autosampler temperature (Ultimate WPS-3000 UHPLC system, Dionex, Germany) was set at 4° C and the injection volume for each sample was 5 µL.

For the chromatographic part (UPLC Ultimate WPS-3000 system Dionex, Germany), we used a C18-XB column (1.7 m, 100 Å, 150×2.1 mm) maintained at 40° C. A mixture of two solvents was used (Solv A: H_2_O + 0.1% formic acid, Solv B: MeOH + 0.1% formic acid) at a flow rate of 0.4 mL/min. The gradient used for the two ionization modes is as follows: 0 to 2 min (A: 99.9%, B: 0.1%); 2-6 min (A: 75%, B: 25%); 6 to 10 min (A: 20%, B: 80%); 10 to 12 min (A: 10%, B: 90%); 12 to 23 min (A: 0.1%, B: 99.9%); 23 to 26.5 min (A: 99.9%, B: 0.1%).

HESI (heated electrospray ionization) source parameters were, for both modes, a spray voltage of 3 kV, capillary temperature of 325° C, heater temperature of 325° C, sheath gas flow of 35 arbitrary units (AU), auxiliary gas flow of 10 AU, sweep gas flow of 1 AU, and S lens RF level of 60 V. During the full-scan acquisition, which ranged from 58 to 870 m/z, the instrument operated at a 70 000 resolution (m/z=200), with an automatic gain control (AGC) target of 1×10^6^ charges and a maximum injection time (IT) of 250 msec. A systematic search for metabolites contained in a library of standard compounds (Mass Spectroscopy Metabolite Library of MSML^®^ Standards, IROA Technologies™) was performed. In order to validate the identity of each detected metabolite, several criteria were required: a) the retention time of the metabolite detected must be within ±20 sec of the standard reference, b) the exact measured molecular exact mass of the metabolite must be within 10 ppm of the known mass of the reference compound, and c) the isotope ratios of the metabolite must match the standard reference. The signal was calculated using Xcalibur^®^ software (Thermo Fisher Scientific, San Jose, CA, USA) by integrating selected ion chromatographic peak area. The data output provides only metabolites for which standard compounds have been validated. The metabolites identified after positive and negative ESI mode analysis were combined to provide a non-redundant list of metabolites useful for statistical analysis. Metabolites with relative standard deviation (RSD) in QCs higher than that in samples were excluded. Only metabolites with RSD in QCs below 30 % and identified in samples were kept for further analysis. Metabolites greater than 30 % variance in QCs were not considered, except if significant variance was observed between groups, meaning that biological variability may exceed analytical variability (59).

### Statistical Analysis

On the evaluation of motor success during single-pulse USNS, data was fit into a sigmoidal curve using a Boltzmann equation (34) of the form:

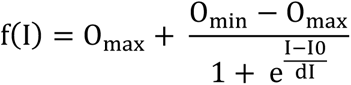

where O_max_ is the highest output (set at 10) and O_min_ the barest (set at 0), I0 is the input half-maximal value and dI the slope.

On the evaluation of the most effective pressure step above the motor threshold, a repeated measure 1-way analysis of variance (ANOVA) was performed with corrected multiple comparisons (post-hoc Tukey). Coefficient of variation (relative standard variation, RSD) were computed for each pressure step in order to characterize high-varying distributions.

On the study of acute repeated USNS and c-Fos densities, repeated measures 2-way ANOVAs were performed accounting for stimulation (acute USNS/Sham) and bregma with corrected multiple comparisons (post-hoc Tukey). The activity index of acute USNS, that is, the relative increase of c-Fos density elicited by acute USNS, was computed to provide effect sizes (difference of means over standard error of the difference).

On the study of chronic USNS, behavioral data was processed with 1-way ANOVAs with corrected multiple comparisons (post-hoc Tukey). Naive mice (non-stressed, untreated) were displayed as ground values and left out of the statistical analysis.

On the study of microPET data, a voxel-based analysis was also used to assess the differences in cerebral [^18^F]-FDG uptake between chronic USNS mice and Sham mice. The ROIs were derived from Mirrione’s templates (58) using PMOD v3.2 software (PMOD Technologies Ltd, Switzerland) and applied to Z-score maps to obtain the Z-score values in these regions. Inter-group comparison was performed using a two-tail unpaired student t-test (XLSTAT). Differences were considered significant when p<0.05

On the study of metabolomics, a first univariate statistical Mann-Withney analysis (XLSTAT) was performed to select metabolites whose expression was significantly different between chronic USNS mice and Sham mice. Next, metabolites whose expression ratios (fold change: FC) between chronic USNS and Sham were greater than 1.25 or less than 0.75 were selected. The pathway enrichment analysis was conducted by the free web software Metaboanalyst (60) to map mouse metabolic pathways corresponding to metabolites selected prior to analysis. The pathway plots were based on the Kyoto Encyclopedia of Genes and Genomes (KEGG) database, and the National Center for Biotechnology Information (NCBI) database was searched to define gene functions. Only the metabolic pathways for which the FDR (false discovery rate) corrected statistics is significant were retained for discussion.

## Acknowledgements

The authors acknowledge the technical platforms (i.e., Department of chemistry and biomedical analyses) of University of Tours, JY Tartu (UMR 1253, iBrain, Tours, France) for designing the stereotaxic manipulator. The work was supported by Inserm Imaging and Brain research unit.

## Authors contributions

M.L., L.G., P.E., C.B. and A.B. designed research; M.L., L.G., A.N., B.P., B.B. and P.G. performed research; B.P. performed c-Fos immunolabelling and analyzed c-Fos data blinded to the conditions; M.L., L.G., S.L., P.E., C.B. and A.B. analyzed data; C.B. and A.B. supervised; M.L., L.G., P.E., C.B., and A.B. wrote paper; M.L., L.G., A.N., B.B., S.L., J.M.E., A.L., P.G., W.E.L., P.E., C.B. and A.B. reviewed paper.

## Supplementary Information

### Supplementary A. Acute repeated USNS evokes neural activity in distant subcortical regions

**Supplementary Figure 1.**
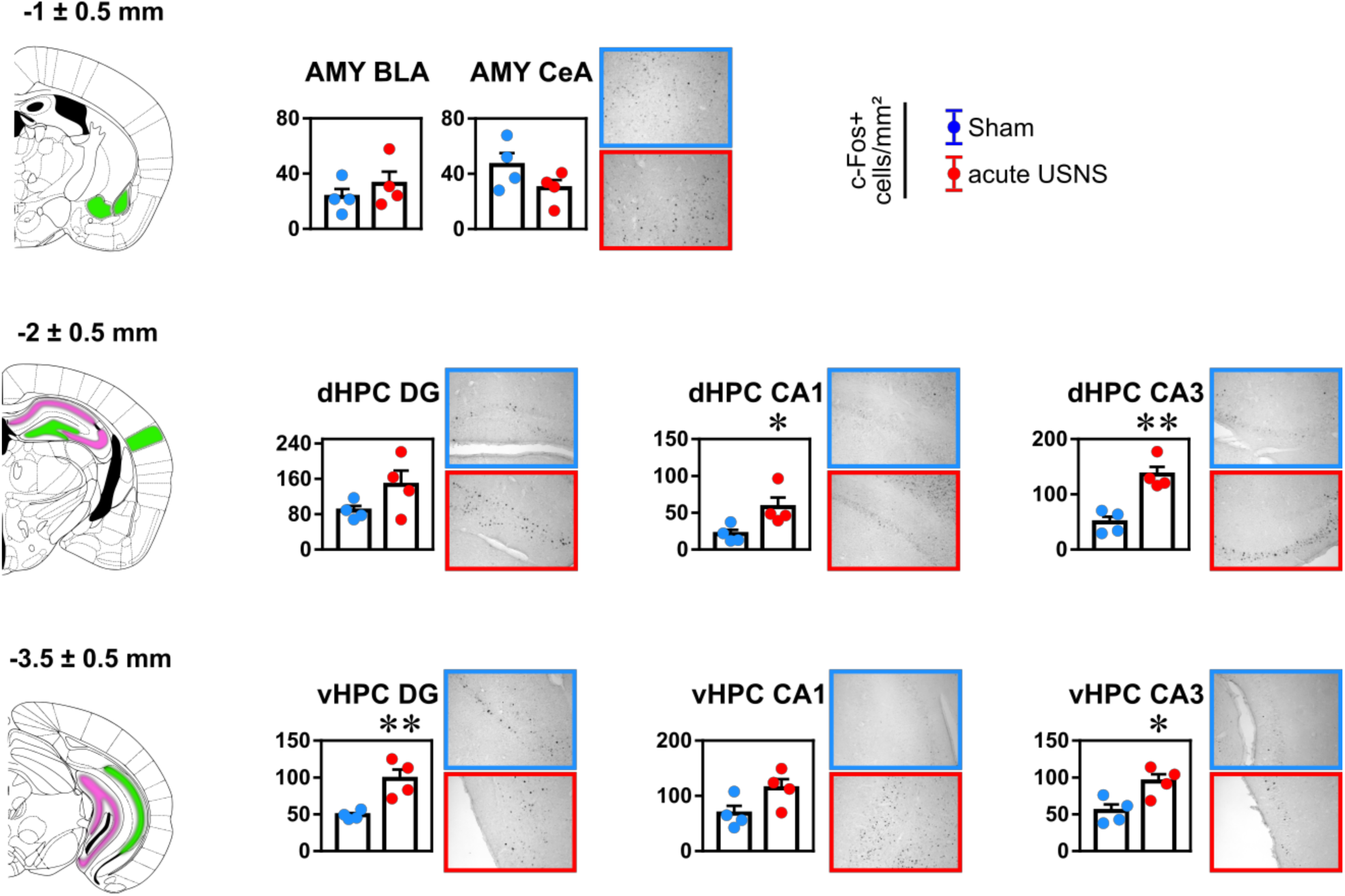
Region locations relative to bregma (mm), c-Fos+ cells/mm² and corresponding micrographs (×20) for (top panel) the amygdala, (mid panel) the dorsal hippocampus and (bottom panel) the ventral hippocampus. Magenta: significant difference, Green: ns. acute USNS: acute repeated USNS, AMY: amygdala, BLA: basolateral amygdala, CeA: central amygdala, HPC: hippocampus, DG: dentate gyrus, CA1, CA3. Data expressed as mean±SEM. * p<0.05, ** p<0.01.

### Supplementary B. Evaluation for hippocampal neurogenesis in chronic USNS mice

On the last day of the experiments (day #43 on the experimental timeline), chronic USNS and Sham mice (n=4/group) were injected with an overdose of pentobarbital (Dolethal) and the brains were harvested and treated as described in Methods, c-Fos analysis. To evaluate the effects of the chronic repeated USNS treatment on neurogenesis, fluorescence IHC was performed on 40-µm thick coronal sections. The doublecortin protein (DCX) was labelled to observe newborn neurons, while the Ki67 protein was labelled to observe proliferating cells. Double-stained cells were taken into account. A classic method was employed, with a primary incubation (1:500 goat anti-DCX, 1:500 rabbit anti-Ki67) at 4° Celsius for 24 hours and a secondary incubation (1:400 donkey anti-goat 555 nm DsRed, 1:400 donkey anti-rabbit 488 nm GFP) at room temperature for 2 hours. Both incubations were done in PB 0.1 M, 2% Normal Donkey Serum and 0.1% Triton and followed by a 15-min washing period in PB 0.1 M (3 × 5 min). The sections were mounted with DAPI (VECTASHIELD Hard Set with DAPI) to label the nucleus of each cell, then observed under a Zeiss Z.2 Imager microscope. For each individual, DCX+, Ki67+ and DCX+/Ki67+ cells were counted in the granular layer of the dentate gyrus in 8 ubiquitous sections along the antero-posterior axis of the brain. Cells counts were expressed as cellular densities (/mm²). For ground values, scores of mice that were treated either with fluoxetine or vehicle were added (the mice followed the same UCMS regimen and displayed similar results and significance in the behavioral tasks described in the main study, n=5/group).

The analysis revealed no difference between chronic USNS and Sham mice (Supplementary Figure 2). Despite an immediate effect of a USNS on c-Fos densities in the hippocampus and the modification of metabolic pathways in the hippocampus following chronic USNS, proliferation mechanisms in the dentate gyrus were not readily engaged in the present case. Because neurogenesis has been associated to antidepressant effects (1), the fact that USNS does not promote the proliferation of newborn neurons could be associated to the lack of effects on depressive-like behaviors at this epoch, measured in the reward-maze test. Independently, mice treated with fluoxetine (Flx) displayed higher DCX+/Ki67+ densities in comparison to vehicle mice (t(8)=4.1, p=0.0035).

**Supplementary Figure 2.**
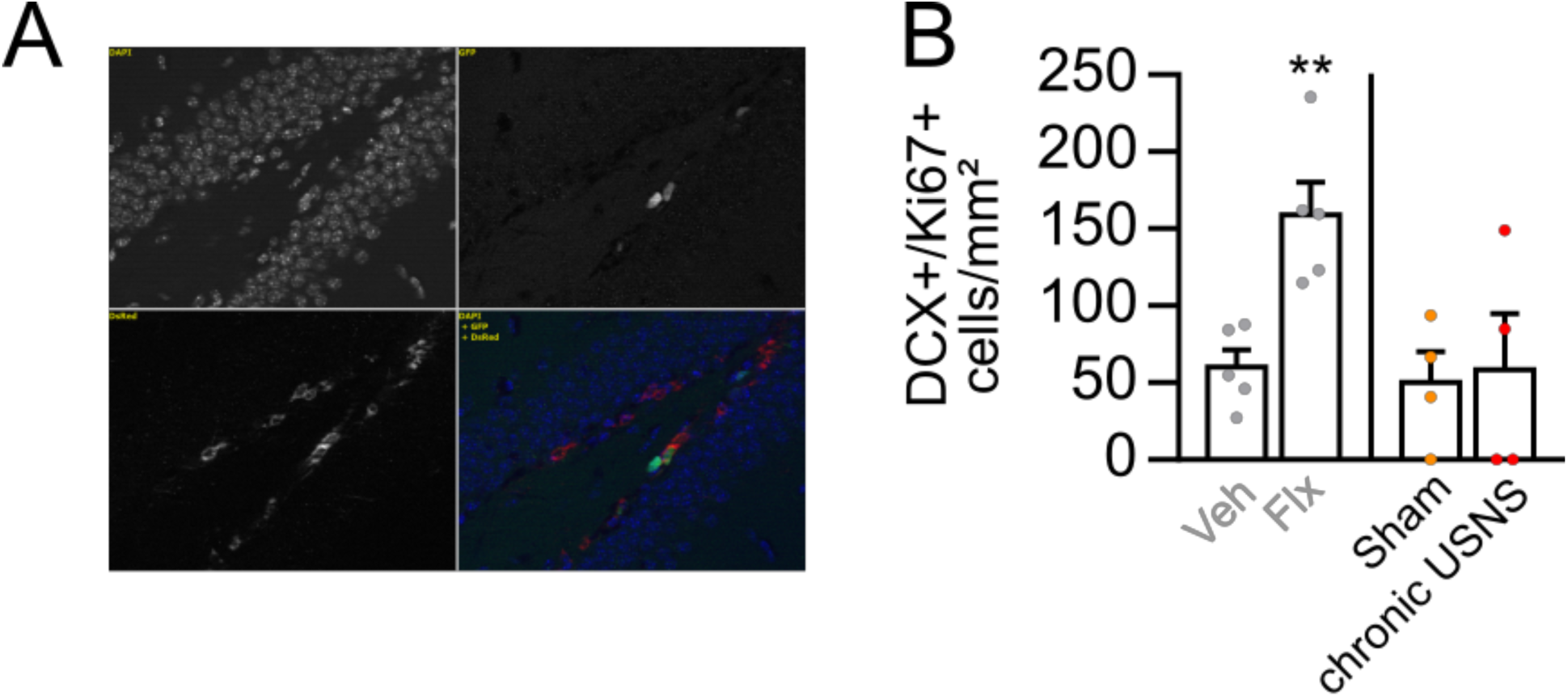
(A) DCX/Ki67 (red/green) double-staining and DAPI (blue). (B) DCX+/Ki67+ cells/mm² for Sham/chronic USNS mice, qualitatively compared to independent values for Veh/Flx mice (in gray). Data expressed as mean±SEM. ** p<0.01.

### Supplementary C. Coat state at day 7

Comparison of coat states with non-stressed mice (NS, n=48), no difference in terms of age or weight from the UCMS mice of the main cohort (n=53). Treatment groups for UCMS (Veh, Flx, Sham, chronic USNS) were determined at day 7 (semi-randomization of mice according to coat states). UCMS deteriorated significantly the coat state at day 7, marking the onset of the UCMS regimen (t(99)=23, p<0.0001; Supplementary Figure 3).

**Supplementary Figure 3.**
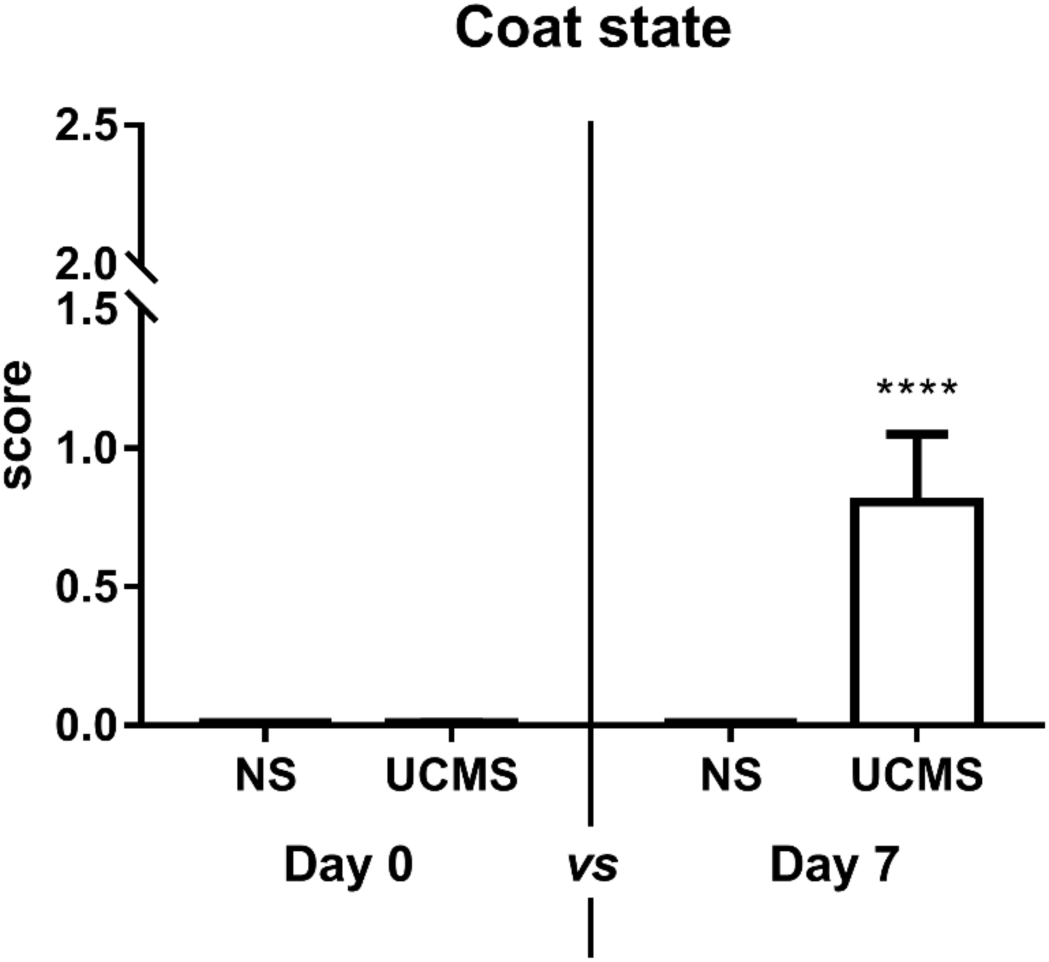
NS: non-stressed, UCMS: unpredictable chronic mild stress. Data expressed as mean ± SEM. **** p<0.0001.

